# Vaccinomics strategy for developing a unique multi-epitope monovalent vaccine against *Marburg marburgvirus*

**DOI:** 10.1101/484923

**Authors:** Mahmudul Hasan, Kazi Faizul Azim, Aklima Begum, Noushin Anika Khan, Tasfia Saiyara Shammi, Md. Abdus Shukur Imran, Ishtiak Malique Chowdhury, Shah Rucksana Akhter Urme

**Author notes:** **Correspondence:** Mahmudul Hasan Assistant Professor Department of Pharmaceuticals and Industrial Biotechnology Faculty of Biotechnology and Genetic Engineering Sylhet Agricultural University, Sylhet-3100, Bangladesh *E-mail:.

## Abstract

Marburg virus causes severe hemorrhagic fever in both humans and non-human primates with high degree of infectivity and lethality. To date no approved treatment is available for Marburg virus infection. A study was employed to design a novel chimeric vaccine against Marburg virus by adopting reverse vaccinology approach. Envelope glycoprotein and matrix protein VP40 were identified as most antigenic viral proteins which generated a plethora of antigenic epitopes. Results showed that vaccine construct V1 was superior in terms of various physicochemical properties and structural stability. Molecular docking analysis of the refined vaccine with different MHCs and human immune TLR8 receptor demonstrated higher binding affinity. Moreover, complexed structure of the modeled vaccine and TLR8 indicated minimal deformability at molecular level. Translational potency and microbial expression of the modeled vaccine within *E. coli* strain K12 by pET28a(+) vector were also biologically significant. However, further *in vitro* and *in vivo* investigation could be implemented for the acceptance and validation of the designed vaccine against Marburg virus.

## 1. Introduction

Marburg virus is the first filovirus ever detected in human. In both humans and nonhuman primates, it causes a severe hemorrhagic fever, known as Marburg hemorrhagic fever (MHF).^1^ The average case lethality rate of Marburg virus disease (MARV) since its first recognition in 1967 is 80%. To date 452 cases and 368 documented deaths have been reported as a result of this zoonotic disease. However, the literature reports suggesting that the actual numbers might be higher.^2,3,4^

Marburg virus belongs to the same virus family to which Ebola virus belongs named filoviridae. The genus *Marburgvirus* includes a single species, *Marburg marburgvirus* first made its appearance in Europe causing severe and fatal hemorrhagic fever among the laboratory workers in Marburg and Frankfurt.^5,6^ About 4 weeks later further cases were observed in Belgrade. Marburg virus reemerged in two large outbreaks in Congo (DRC) in 1998–2000 and then, for the first time in West African country Angola in 2004– 2005.^7,8^ At that time, a total 406 cases were observed with a high fatality rate of 83% and 90% in Congo and Angola respectively. Those incidences revealed MARV as major threat for public health.^8,9,10^

Three Marburg virus disease outbreaks have been reported in Uganda with first one documented in 2007. In 2012, MHF was responsible for with 15 deaths among 26 cases observed in multiple districts.^11^ Close interaction between people and animals such as non-human primates, bats, and livestock was attributed as a probable cause of those outbreaks. Egyptian fruit bat (Rousettus aegyptiacus) is currently known as a reservoir of Marburg viruses^12^ and cases have been linked to exposure in caves or mines residing by this organism.^13,14^ The pathological symptoms of the disease involve a number of systemic dysfunctions including edema, hemorrhages, shock, multi organ failure, often resulting in death.^15^

Due to its high infectivity and lethality, handling of Marburg virus is restricted to high containment biosafety Level 4 laboratories.^16^ It has also been classified as category A priority pathogen by the National Institute of Allergy and Infectious Diseases (NIAID), select agent by the Centers for Disease Control and Prevention (CDC) and risk group 4 agent by WHO. Currently, there are no vaccines or drugs approved for human to protect against Marburg virus.^2^ Supportive care (fluids, antimicrobials, blood transfusion) has been the primary treatment of patients during the outbreaks.^3,4^

The conventional approach for vaccine designing is time consuming and only allows the identification of abundant antigens that are cultivable under laboratory conditions. Initial approaches using inactivated virus to develop a vaccine against MARV were unsuccessful or had contradictory results.^17^ But, the reverse approach to vaccine development overcomes these problems taking benefit of the genome sequence of the pathogen. The strategy aims to combine immunogenomics and immunogenetics with bioinformatics for the development of novel vaccine target.^18^ Subunit vaccines consist of only the antigenic part of the pathogen with the potential to induce a protective immune response inside host while overcoming the problems caused by live attenuated vaccines.^19^

Marburg virus possesses a single surface protein named envelope glycoprotein (GP) that mediates attachment to target cells and virus entry.^20^ Besides its function in entry, GP is also associated with immune evasion. Another protein of this virus matrix protein (VP40) plays a major role in the formation of virions and recruiting GP.^21^ Both the proteins hold potential to be an effective target for designing a vaccine against Marburgvirus. The present study was conducted to design a unique, non-allergic and immunogenic chimeric vaccine against Marburgvirus utilizing the vaccinomics approach while the wet lab researchers are anticipated to authenticate our prediction.

## 2. Materials and methods

In the present study, an *in silico* approach was empoyed to design vaccine candidates against Marburgvirus to prevent MHF. A flow chart showing the protocol over vaccinomics approach for developing an epitope-based chimeric vaccine has been illustrated in figure 1.

**Fig. 1.**
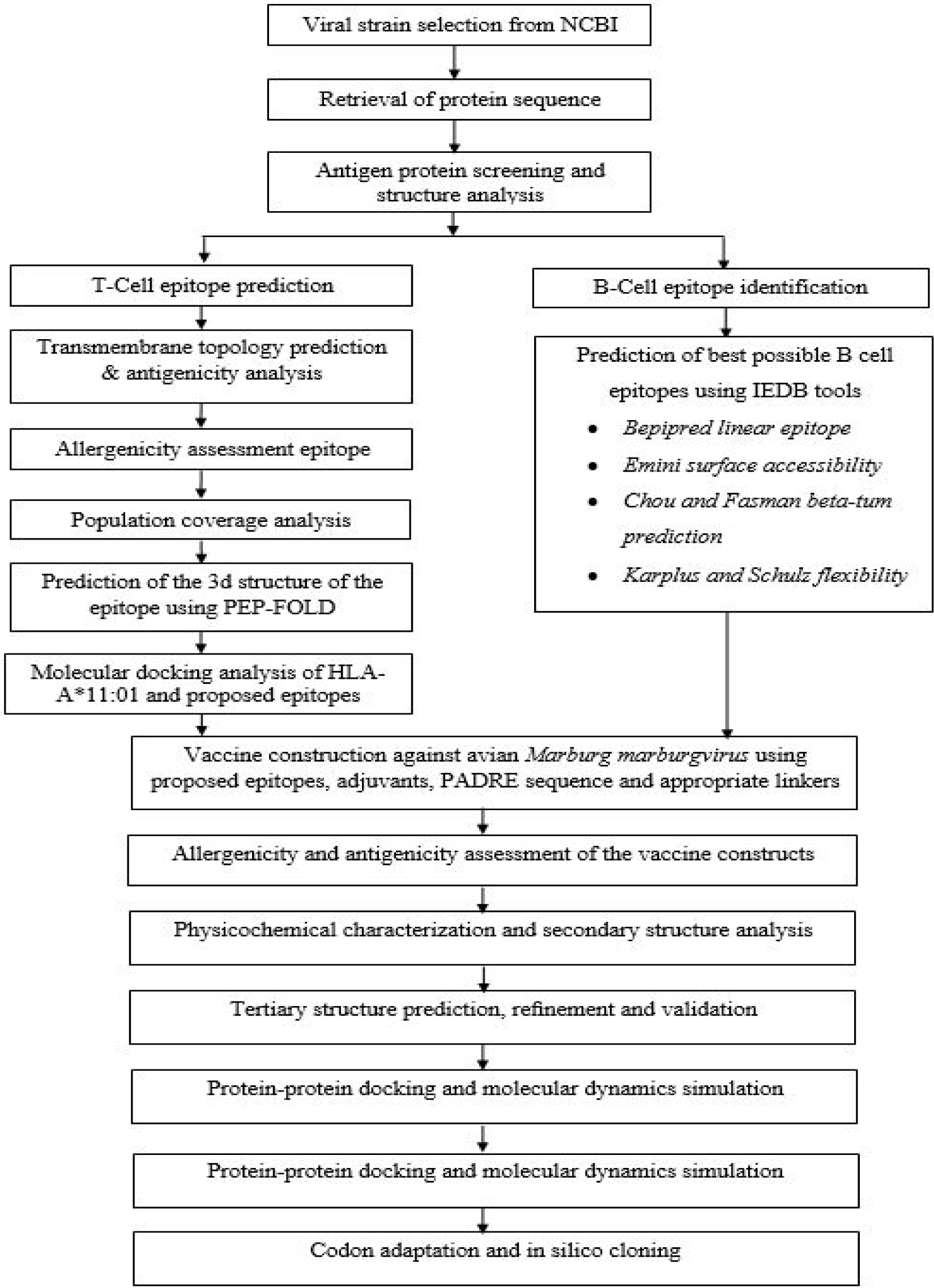
Flow chart summarizing the protocols for the prediction of epitope based vaccine candidate by *in silico* reverse vaccinology technique.

### 2.1. Viral strain selection

The National Center for Biotechnology Information (NCBI) was used for the selection of Marburgvirus *(Marburg marburgvirus)* strain Musoke-80 (https://www.ncbi.nlm.nih.gov/genome/genomes/5). The server provides access to biomedical and genomic information over numerous organisms. Study of other associated information including the genus, family, host, transmission, disease, genome and proteome analysis were performed by using ViralZone, a web-resource of Swiss Institute of Bioinformatics (https://viralzone.expasy.org/194).

### 2.2. Protein sequence retrieval

UniProt is database containing a large amount of information about the biological information of protein. The entire viral proteome of Marburgvirus *(Marburg marburgvirus)* strain Musoke-80 was retrieved from UniProtKB (https://www.uniprot.org/uniprot/?query+database) consisting seven proteins.

### 2.3. Antigenic protein screening and structure analysis

Antigenicity refers to the capacity of antigens to be recognized by the immune system. To investigate protein antigenicity and determine the most potent antigenic proteins, the VaxiJen v2.0 server (http://www.ddg-pharmfac.net/vaxijen/) was utilized^22^. From the seven viral proteins, the two structural proteins (envelope glycoprotein and matrix protein VP40) were selected based on their antigenic score. Different physicochemical properties of the proteins were predicted using ProtParam^23^, one of the ExPASy’s servers for primary structure prediction of proteins.

### 2.4. T-Cell epitope prediction

The IEDB offers easy searching of experimental data characterizing antibody and T cell epitopes studied in human and other non-human primates. From this Immune Epitope Database, MHC-I prediction tool (http://tools.iedb.org/mhci/) and MHC-II prediction tool (http://tools.iedb.org/mhcii/) were used to predict the MHC-I binding and MHC-II binding respectively.^24^ Both MHC-I restricted CD8+ cytotoxic T lymphocytes (CTLs) and MHC-II restricted CD4+ cytotoxic T lymphocytes play a pivotal role in controlling viral infections. Hence, identification of T cell epitopes is crucial for understanding the mechanism of T cell activation and epitope driven vaccine design. The protein sequences, envelope glycoprotein and matrix protein VP40 were added to the query box at different time for both type of analysis. MHC class I and MHC class II alleles were selected.

### 2.5. Transmembrane topology prediction and antigenicity analysis of epitopes

The TMHMM server (http://www.cbs.dtu.dk/services/TMHMM/) predicted the transmembrane helices in proteins. The topology was determined according to the position of the transmembrane helices separated by ‘i’ if the loop is on the inside or ‘o’ if it is on the outside.^25^ Again, VaxiJen v2.0 server (http://www.ddg-pharmfac.net/vaxijen/) was used to determine the epitope antigenicity.^22^ The most potent antigenic epitopes were selected for further investigation.

### 2.6. Population coverage analysis

HLA distribution varies among different ethnic groups and geographic regions around the world. So, population coverage must be taken into account when designing an effective vaccine to cover as much as possible populations. In this study, population coverage for each individual epitope was analyzed by the IEDB population coverage calculation tool analysis resource (http://tools.iedb.org/population/).

### 2.7. Allergenicity assessment and toxicity analysis of T-Cell epitopes

The prediction of allergens has been explored widely using bioinformatics, with many tools being developed in the last decade. Four servers named AllerTOP (http://www.ddg-pharmfac.net/AllerTop/),^26^ AllergenFP (http://www.ddg-pharmfac.net/AllergenFP/),^27^ PA^3^P (http://www.Ipa.saogabriel.unipampa.edu.br:8080/pa3p/)^28^ and Allermatch (http://www.allermatch.org/allermatch.py/form)^29^ were used to predict the allergenicity of our proposed epitopes for vaccine development. Only the non-allergenic epitopes were allowed to demonstrate the toxicity level by ToxinPred server (http://crdd.osdd.net/raghava/toxinpred/).

### 2.8. Conservancy analysis

Epitope conservancy is a vital step in the immunoinformatic approach as it determines the extent of desired epitope distributions in the homologous protein set. IEDB’s epitope conservancy analysis tool (http://tools.iedb.org/conservancy/) was selected for the analysis of conservancy level by concentrating on the identities of the selected proteins.

### 2.9. Cluster analysis of the MHC restricted alleles

Structure based clustering methods have been proven efficient to identify the super-families of MHC proteins with similar binding specificities. Due to the vast polymorphic nature among species, the distinct specificity of the MHC alleles remains uncharacterized in most cases. In this study, a tool from MHCcluster v2.0^30^ server was used to produce pictorial tree-based visualizations and highly instinctive heat-map of the functional alliance between MHC variants.

### 2.10. Designing three-dimensional (3D) epitope structure

The top ranked epitopes were subjected for the docking study after analyzing through different bioinformatics tools. PEP-FOLD is a *de novo* approach aimed at predicting peptide structures from amino acid sequences.^31^ By Folding peptides on a user specified patch of a protein, it comes with the possibility to generate candidate conformations of peptide-protein complexes^32^. The server was used to design and retrieve the 3D structure of most potent selected epitopes for further analysis.

### 2.11. Molecular docking analysis

MGLTools is a software developed for the visualization and analysis of molecular structures.^33^ It includes AutoDock, an automated docking software designed to predict the interactions between small molecules (i.e. substrates or drug candidates) and receptor of known 3D structure.^34^ All the operations were performed at 1.00°A space keeping the exhaustiveness parameter at 8.00. The numbers of outputs were set at 10. The docking was conducted using AutoDOCK Vina program based on the above-mentioned parameters. OpenBabel (version 2.3.1) was used to convert the output PDBQT files in PDB format. The best output was selected on the basis of higher binding energy. The docking interaction was visualized with the PyMOL molecular graphics system, version 1.5.0.4 (https://www.pymol.org/).

### 2.12. B-Cell epitope identification

The objective of B cell epitope prediction was to find the potential antigen that would interact efficiently with B lymphocytes and initiate an immune response. From the experimental confirmation, it was confirmed that the flexibility of the peptide is associated to its antigenicity. For a B cell to be potential epitope, it must have proper surface accessibility as well. B cell epitope prediction tools from IEDB were used to identify the B cell antigenicity based on six different algorithms which include Kolaskar and Tongaonkar antigenicity scale,^35^ Emini surface accessibility prediction,^36^ Karplus and Schulz flexibility prediction,^37^ Bepipred linear epitope prediction analysis,^38^ Chou and Fasman beta turn prediction^39^ and Parker hydrophilicity prediction.^40^

### 2.13. Vaccine construction

Subunit vaccines consist of antigenic parts of a pathogen to stimulate an immunogenic reaction in the host. The predicted T-cell and B-cell epitopes were conjugated in a sequential manner to design the final vaccine construct. All three vaccine proteins started with an adjuvant followed by the top CTL epitopes for both capsid protein VP1 and protein VP2, then by top HTL epitopes and BCL epitopes respectively, in the similar fashion. Three vaccine sequence were constructed named V1, V2 and V3, each associated with different adjuvants including beta defensin (a 45 mer peptide), L7/L12 ribosomal protein and HABA protein (*M. tuberculosis*, accession number: AGV15514.1). Interactions of adjuvants with toll like receptors (TLRs) polarize CTL responses and induce robust immunoreaction.^41^ Beta defensin adjuvant can act as an agonist to TLR1, TLR2 and TLR4 receptor. On the contrary, L7/L12 ribosomal protein and HBHA protein are agonists to TLR4 only. To overcome the problems caused by highly polymorphic HLA alleles, PADRE sequence was also incorporated along with the adjuvant peptides. EAAAK linkers were used to join the adjuvant and CTL epitopes. Similarly, GGGS, GPGPG and KK linkers were used to conjugate the CTL, HTL and BCL epitopes respectively. Utilized linkers ensured the effective separation of individual epitopes in vivo.^42,43^

### 2.14. Allergenicity and antigenicity prediction of different vaccine constructs

AlgPred v.2.0^29^ sever was used to predict the non-allergic nature of the constructed vaccines. The server developed an algorithm by considering the auto cross covariance transformation of proteins into uniform vectors of similar length. The accuracy of results ranges from 70% to 89% depending on species. We further used VaxiJen v2.0 server^22^ to evaluate the probable antigenicity of the vaccine constructs in order to suggest the superior vaccine candidate. The server analyzed the immunogenic potential of the given proteins through an alignment-independent algorithm.

### 2.15. Physicochemical characterization and secondary structure analysis of vaccine protein

ProtParam, a tool provided by Expasy server^23^ was used to functionally characterize the vaccine constructs according to molecular weight, aliphatic index, isoelectric pH, hydropathicity, instability index, GRAVY values, estimated half-life and various physicochemical properties. By comparing the pK values of diferent aminoacids, the server computes these parameters of a given protein sequence. Aliphaticindex is the volume occupied by the aliphatic side chains of protein. Grand average of hydropathicity was computed by summing the hydropathicity of all amino acid residues present in the protein sequence and then by dividing it by total number of amino acid residues. The PSIPRED v3.3^44^ and NetTurnP 1.0 program^45,46^ was used to predict the alpha helix, beta sheet and coil structure of the vaccine constructs.

### 2.16. Vaccine tertiary structure prediction, refinement and validation

The RaptorX server performed 3D modeling of the designed vaccines depending on the degree of similarity between target protein and available template structure from PDB. 47,48 Refinement was conducted using ModRefiner^49^ followed by FG-MD refinement server^50^ to improve the accuracy of the predicted 3D modeled structure. ModRefiner drew the initial model closer to its native state based on hydrogen bonds, side-chain positioning and backbone topology, thus resulting in significant improvement in the physical quality of the local structure. FG-MD is another molecular dynamics based algorithm for structure refinement at atomic level. The refined protein structure was further validated by Ramachandran plot assessment at RAMPAGE.^51^

### 2.17. Vaccine protein disulfide engineering

Disulfide bonds enhance the geometric conformation of proteins and provide significant stability. DbD2, an online tool was used to design such bonds for the constructed vaccine protein.^52^ The server detects and provides a list of residue pairs with proper geometry which have the capacity to form disulfide bond when individual amino acids are mutated to cysteine.

### 2.18. Protein-protein docking

Molecular docking aims to determine the binding affinity between a receptor molecule and ligand.^53^ Inflammations caused by single stranded RNA virus are involved with immune receptors, mainly by TLR-7 and TLR-8 present over the immune cells.^54,55^ An approach for protein-protein docking was employed to determine the binding affinity of designed subunit vaccines with different HLA alleles and TLR-8 immune receptor by using ClusPro 2.0.^56^, hdoc^57,58^ and PatchDock server.^59^ The 3D structure of different MHC molecules and human TLR-8 receptor was retrieved from RCSB protein data bank. The above mentioned servers were used to obtain the desirable complexes in terms of better electrostatic interaction and free binding energy. PatchDock generated a number of solutions which were again subjected to the FireDock server to refine the complexes.

### 2.19. Molecular dynamics simulation

Molecular dynamics study is important to strengthen any *in silico* prediction and demonstrate the stability of protein-protein complex. Stability can be determined by comparing the essential dynamics of proteins to their normal modes.^60,61^ This powerful tool is an alternative to the costly atomistic simulation.^62,63^ iMODS server explains the collective motion of proteins by analyzing the normal modes (NMA) in internal coordinates.^64^ The structural dynamics of protein complex was investigated by using this server due to its much faster and effective assessments than other molecular dynamics (MD) simulations tools.^65,66^ It predicted the direction and extent of the immanent motions of the complex in terms of deformability, eigenvalues, B-factors and covariance. The deformability of the main chain depends on the ability to deform at each of its residues for a given molecule. The eigenvalue related to each normal mode describes the motion stiffness. This value is directly linked to the energy required to deform the structure. Deformation is much easier if the eigenvalue is low and vice versa.^67^

### 2.20. Codon adaptation and in silico cloning

Codon adaptation tools are used for adapting the codon usage to the well characterized prokaryotic organisms to accelerate the expression rate in them. *E. coli* strain K12 was selected as host for cloning purpose of the designed vaccine construct. Due to the lack of similarities between the codon usage of human and *E. coli*, the approach was adopted to achieve higher expression of vaccine protein V1 in the selected host. Rho independent transcription termination, prokaryote ribosome-binding site and cleavage sites of several restriction enzymes (i.e. BglII and Apa1) were avoided during the operation performed by JCAT server.^68^ The optimized sequence of vaccine protein V1 was reversed and then conjugated with BglII and Apa1 restriction site at the N-terminal and C-terminal sites respectively. SnapGene^53^ restriction cloning module was used to insert the adapted sequence between BglII (401) and ApaI (1334) of pET28a(+) vector.

## 3. Results

### 3.1. Protein sequence retrieval

The entire viral proteome of Marburgvirus *(Marburg marburgvirus)* Musoke-80 strain was extrated from UniProtKB (https://www.uniprot.org/uniprot/?query+database). All the sequences were from Kenya. The viral proteome consists seven viral proteins named envelope glycoprotein, matrix protein VP40, RNA directed RNA polymerase L, nucloeoprotein, polymerase cofactor VP35, membrane-associated protein VP24 and minor nucleoprotein VP30.

### 3.2. Antigenic protein prediction and structure analysis

The VaxiJen server was used to assess the retrieved protein sequences in order to find the most potent antigenic protein. Envelope glycoprotein (Accession ID: P35253) and matrix protein VP40 (Accession ID: P35260) were selected as the most potent antigenic protein with total prediction score of 0.5474 and 0.4107 respectively and allowed for further analysis. Various physiochemical parameters of the proteins were analyzed by ProtParam tools as listed in table 1.

**Table 1:** ProtParam analysis of retrieved viral proteins.

| Proteins | Accession ID | Molecular Weight | Instability Index | Half-life | Theoretical pI | No. of Amino acids | Total No. of Atoms | Extinction Co-efficient |
| --- | --- | --- | --- | --- | --- | --- | --- | --- |
| Envelope glycoprotein | P35253 | 74376.19 | 45.52 | 30 hours | 5.88 | 681 | 10350 | 80620 |
| Matrix protein VP40 | P35260 | 33793.72 | 25.46 | 30 hours | 9.40 | 303 | 4757 | 28880 |

### 3.3. T-Cell epitope prediction

The results were investigated according to epitope-HLA cell binding. Epitopes that bind to the maximum number of HLA cells were selected. Herein, numerous immunogenic epitopes from envelope glycoprotein and matrix protein VP40 were identified to be T cell epitopes using both MHC-I and MHC-II binding predictions of the IEDB that can bind a large number of different HLA-A and HLA-B alleles with high binding affinity.

### 3.4. Transmembrane topology prediction and antigenicity analysis

Top epitopes of both envelope glycoprotein and matrix protein VP40 were selected as putative T cell epitope candidates based on their transmembrane topology screening by TMHMM and antigenic scoring (AS) by Vaxijen (Table 2 and 3). Epitopes with a positive score of immunogenicity exhibited potential to elicit effective T-cell response.

**Table 2:**
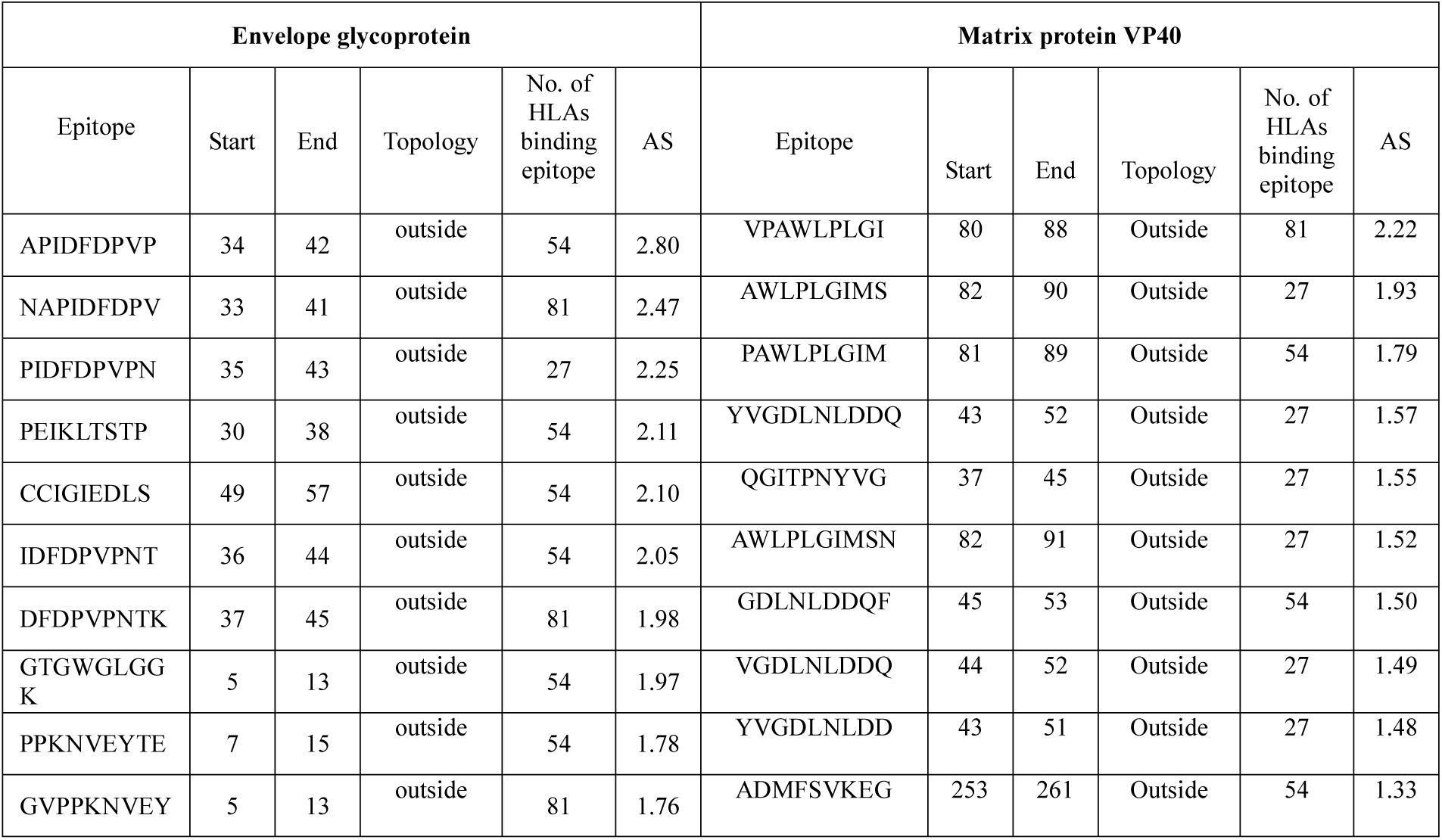
Predicted T-cell epitopes (MHC-I peptides) of envelope glycoprotein and matrix protein VP40.

**Table 3:**
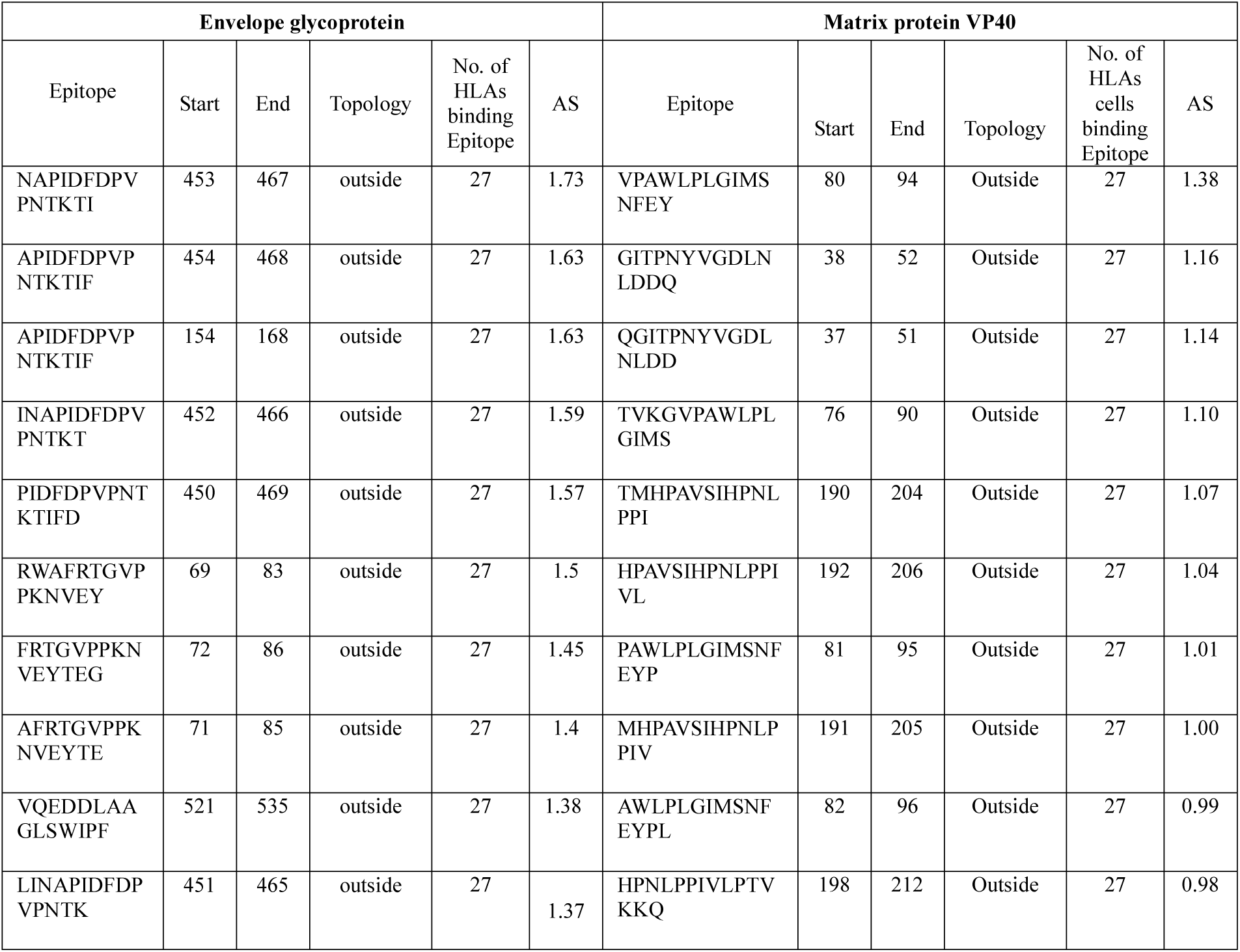
Predicted T-cell epitopes (MHC-II peptides) of envelope glycoprotein and matrix protein VP40.

### 3.5. Population coverage analysis

Population coverage for each individual epitopes was analyzed by the IEDB population coverage calculation tool analysis resource. All indicated alleles in supplementary data were identified as optimum binders with the predicted epitopes and were used to determine the population coverage. Population coverage result for both, envelope glycoprotein and matrix protein VP40 are shown in figure 2.

**Fig. 2.**
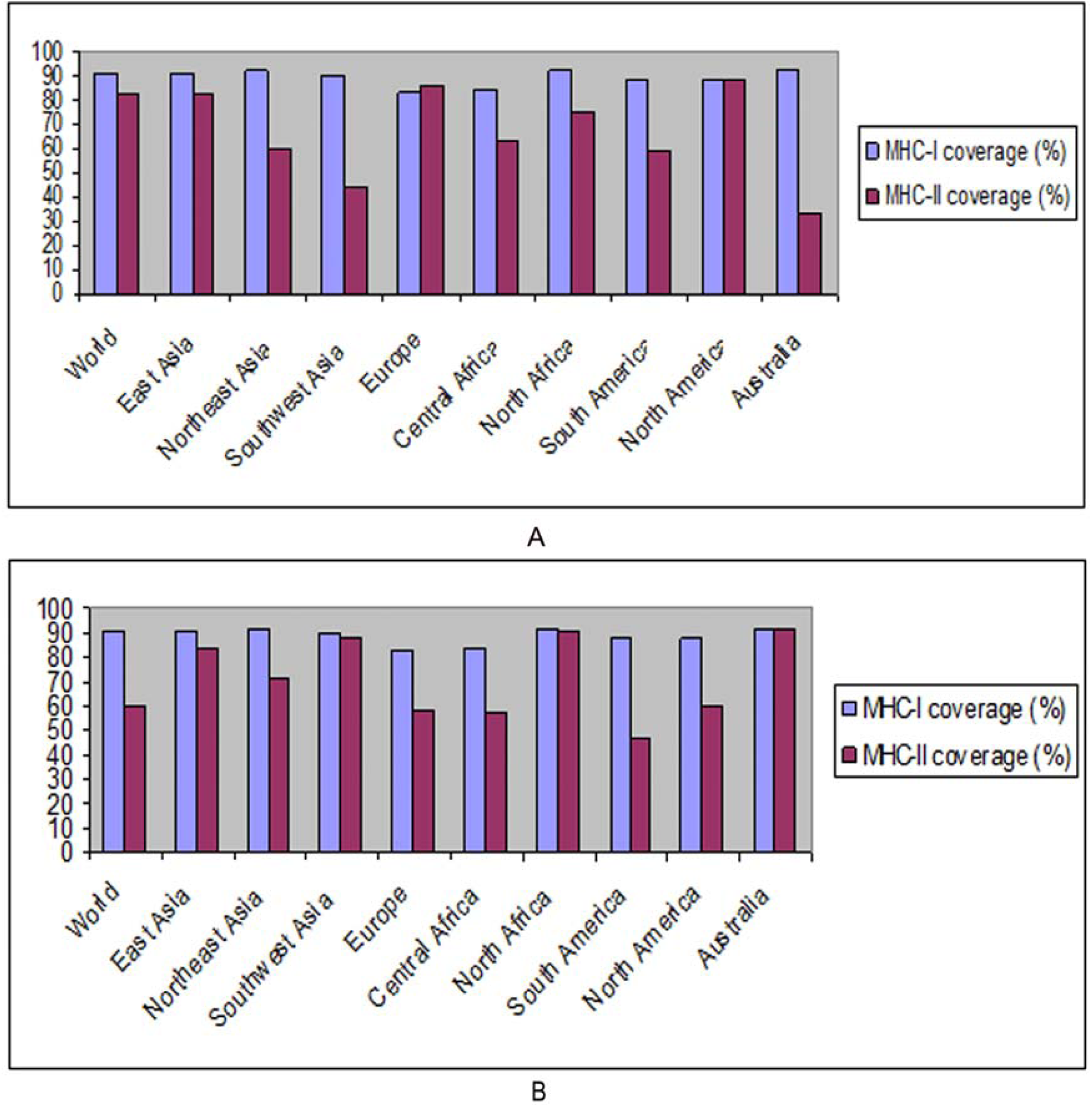
Population coverage analysis of envelope glycoprotein **(A)** and matrix protein VP40 **(B).**

### 3.6. Allergenicity assessment and toxicity analysis of T-Cell epitopes

Based on the allergenicity assessment by four servers (i.e. AllerTOP, AllergenFP, PA^3^P, Allermatch), epitopes that were found to be non-allergen for human were identified. Epitopes those were indicated as allergenic for human or classified as undefined were removed from the predicted list of epitopes. Epitopes those were predicted as toxic by ToxinPred server were screened out too (Table 4).

**Table 4:** Allergenicity assessment, toxicity test and conservancy analysis of the predicted epitopes generated from envelope glycoprotein and matrix protein VP40.

| Envelope glycoprotein |  |  |  | Matrix protein VP40 |  |  |  |
| --- | --- | --- | --- | --- | --- | --- | --- |
| Epitope | Allergenicity | Toxicity | Conservancy | Epitope | Allergenicity | Toxicity | Conservancy |
| APIDFDPVP | Non Allergen | Non Toxin | 72.9% | VPAWLPLGI | Non Allergen | Non Toxin | 100% |
| CCIGIEDLS | Non Allergen | Toxin | 91.6% | YVGDLNLDD | Non Allergen | Non Toxin | 95.6% |
| GVPPKNVEY | Non Allergen | Non Toxin | 95.8% | QGITPNYVG | Non Allergen | Non Toxin | 52.1% |
| PPKNVEYTE | Non Allergen | Non Toxin | 95.8% | VGDLNLDDQ | Non Allergen | Non Toxin | 95.6% |
| PEIKLTSTP | Non Allergen | Non Toxin | 4.17% | ADMFSVKEG | Non Allergen | Non Toxin | 95.6% |
| NAPIDFDPVP<br>NTKTI | Non Allergen | Non Toxin | 70.8% | VPAWLPLGIM<br>SNFEY | Non Allergen | Non Toxin | 100.% |
| APIDFDPVPN<br>TKTIF | Non Allergen | Non Toxin | 70.8% | GITPNYVGDL<br>NLDDQ | Non Allergen | Non Toxin | 95.6% |
| RWAFRTGVP<br>PKNVEY | Non Allergen | Non Toxin | 89.5% | QGITPNYVG<br>DLNLDD | Non Allergen | Non Toxin | 52.1% |
| VQEDDLAAG<br>LSWIPF | Non Allergen | Non Toxin | 89.5% | TMHPAVSIHP<br>NLPPI | Non Allergen | Toxin | 39.1% |
| FRTGVPPKN<br>VEYTEG | Non Allergen | Non Toxin | 89.5% | HPAVSIHPNL<br>PPIVL | Non Allergen | Non Toxin | 39.1% |

### 3.7. Conservancy analysis

To ensure a broad spectrum immune response by a vaccine, the conservancy level for the epitope candidates must be satisfactory. Here, putative epitopes generated from envelope glycoprotein and matrix protein VP40 were found to be highly conserved with maximum conservancy level of 95.8% and 100% respectively (Table 4).

### 3.8. Cluster analysis of the MHC restricted alleles

MHCcluster v2.0 produced a clustering of both class I and class II HLA molecules which were detected to interact with the predicted epitopes. The result was generated through conventional phylogenetic method based on sequence data available for different HLA-A and HLA-B alleles. Figure 3 is illustrating the function based clustering of HLA alleles (heat map) where red zones indicating strong correlation and yellow zone showing weaker interaction.

**Fig. 3.**
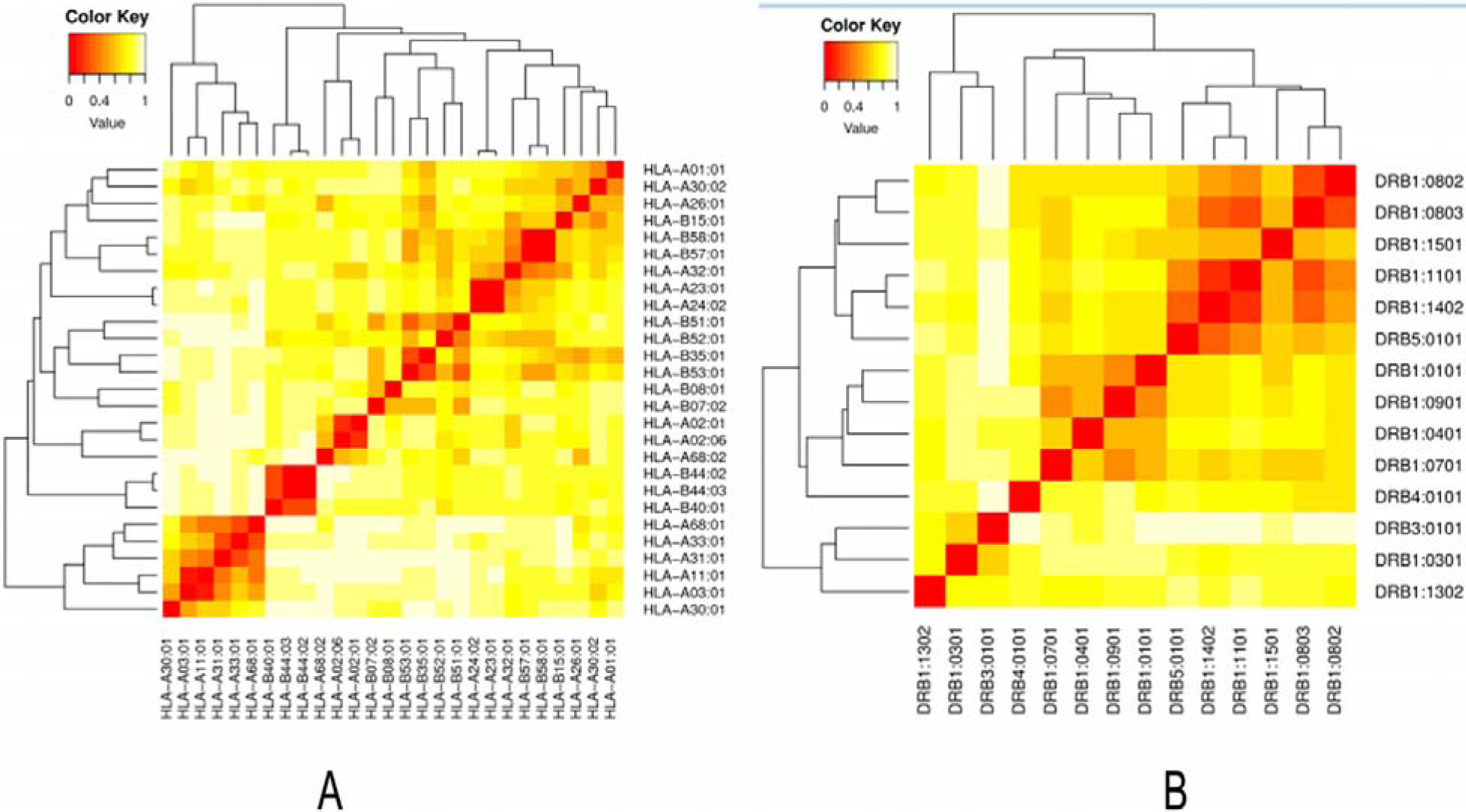
Cluster analysis of the HLA alleles: (**A:** MHC-I molecules, **B:** MHC-II molecules (red color in the heat map indicating strong interaction, while the yellow zone indicating the weaker interaction).

### 3.9. Molecular docking analysis and HLA allele interaction

For docking analysis, 12 T-cell epitopes (six from envelope glycoprotein and six from matrix protein VP40) were subjected to PEP-FOLD3 web-based server for 3D structure conversion in order to analyze their interactions with HLA molecule. From five structures, modeled by the server for each individual epitopes, the best one was identified for docking study. The binding patterns of the HLA molecules and predicted epitopes were investigated using MGLTools. HLA-A*11:01 and HLA-DRB1*04:01 was selected for docking analysis with MHC class I and class II binding epitopes respectively, on the basis of the available Protein Data Bank (PDB) structure deposited in the database. All the selected epitopes were allowed for docking analysis using AutoDock and the binding energies were analyzed (Table 5).

**Table 5:** Binding energy of suggested T-cell epitopes with selected class I and class II MHC molecules generated from molecular docking analysis

| *Epitopes* | *MHC allele* | *Binding Energy (kcal/mol)* | *Epitopes* | *MHC allele* | *Binding Energy (kcal/mol)* |
| --- | --- | --- | --- | --- | --- |
| APIDFDPVP | MHC-A*11:01 | -9.5 | NAPIDFDPVPNTKTI | HLA-DRB1*04:01 | -7.6 |
| GVPPKNVEY |  | -8.8 | RWAFRTGVPPKNVEY |  | -7.5 |
| PPKNVEYTE |  | -7.8 | VQEDDLAAGLSWIPF |  | -7.8 |
| VPAWLPLGI |  | -9.1 | VPAWLPLGIMSNFEY |  | -7.0 |
| YVGDLNLDD |  | -8.6 | GITPNYVGDLNLDDQ |  | -6.5 |
| QGITPNYVG |  | -7.9 | QGITPNYVGDLNLDD |  | -6.2 |

Result showed that ‘VQEDDLAAGLSWIPF’ epitope of envelope glycoprotein (GP) bound in the groove of the HLA-DRB1*04:01 with an energy of -7.8 kcal/mol. The demonstrated energy was -7.0 kcal/mol for epitope ‘VPAWLPLGIMSNFEY’ contained the 9-mer core ‘VPAWLPLGI’ of matrix protein VP40. On the contrary, VP1-epitope ‘APIDFDPVP’ was found to be superior in terms free binding energy while interacted with HLA-A*11:01 (-9.5 kcal/mol). The docked complexes were visualized by the latest version of PyMOL molecular graphics system.

### 3.10. B-Cell epitope identification

B-cell epitopes of both, envelope glycoprotein and matrix protein VP40 were generated using six different algorithms. For envelope glycoprotein, Bepipred prediction method predicted the peptide sequences from 466-489 and 612-658 amino acids as potential B cell epitopes that could induce the preferred immune responses (Figure 4:A). Emini-surface accessibility prediction was also conducted which indicated 293-31 and 626-634 amino acid residues to be more accessible (Figure 4:B). Chou and Fasman beta-turn prediction method displayed regions from 26-32 and 308-314 as potential Beta-turn regions (Figure 4:C). Karplus and Schulz flexibility prediction method found the region of 190-196 and 488-494 amino acid residues as most flexible regions (Figure 4:D). In contrast, Kolaskar and Tongaonkar antigenicity result confirmed the region from 424-432 and 601-614 as highly antigenic (Figure 4:E) while Parker hydrophilicity prediction indicated 407-413 and 606-612 amino acid residues to be more potent (Figure 4:F).

**Fig. 4.**
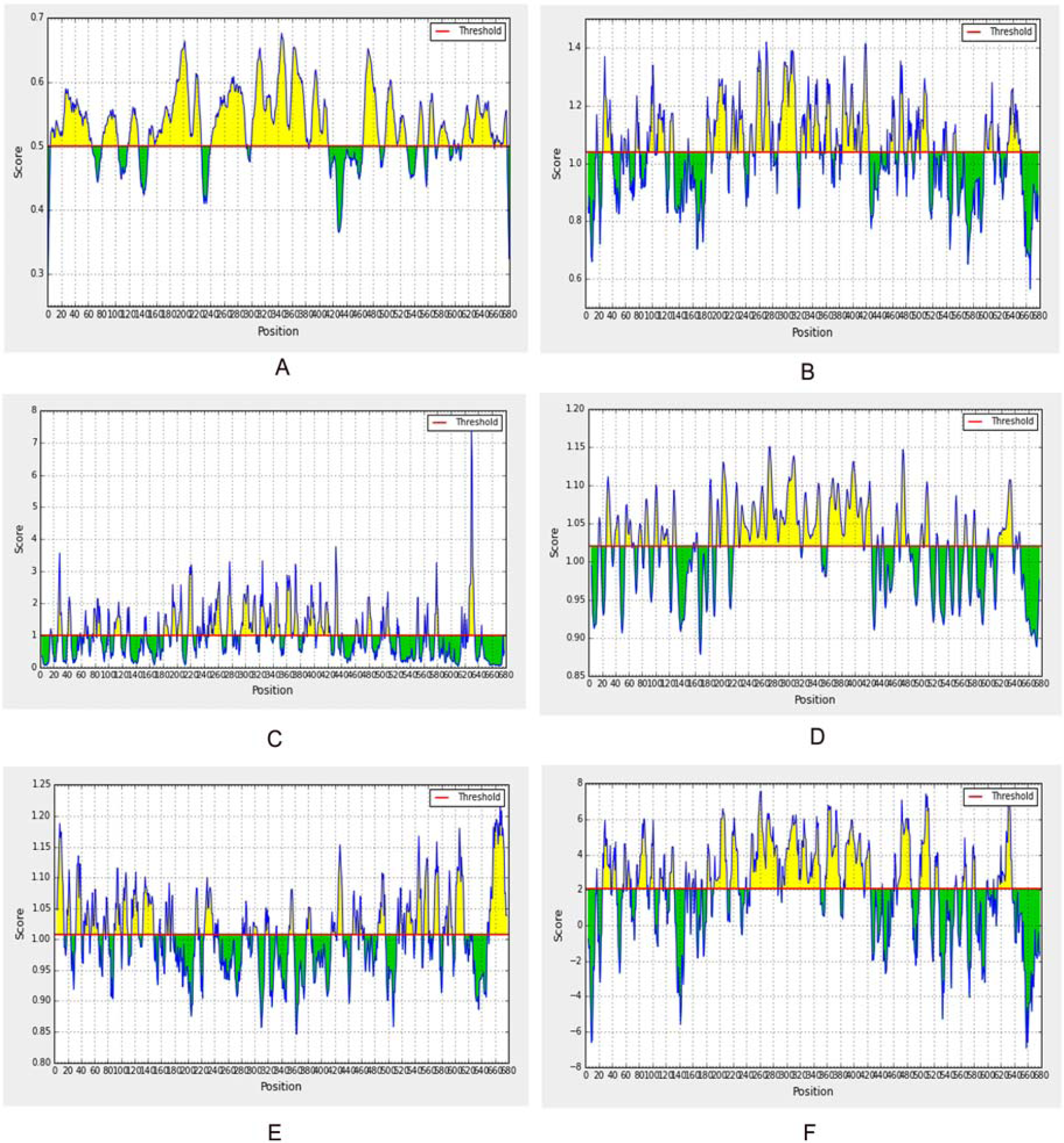
Prediction of B cell linear epitope and intrinsic properties for membrane glycoprotein using different scales (**A:** Bepipred, **B:** Surface accessibility, **C:** Emini surface, **D:** Flexibility, **E:** Antigenicity, **F:** Hydrophilicity). **Notes:** For each graph: x-axis and y-axis represent the position and score; residues that fall above the threshold value are shown in yellow color; the highest peak in yellow color identifies most favored position).

In case of matrix protein VP40, bepipred prediction method identified the peptide sequences from 29-40 and 214-223 amino acids with ability to induce preferred immunity (Figure 5:A). The regions from 177-184 and 208–222 amino acid residues were more accessible based on Emini surface accessibility prediction algorithm (Figure 5:B). Regions from 29–35 and 217–223 were potential beta-turn regions on the basis of Chou and Fasman beta-turn prediction (Figure 5:C). Karplus and Schulz flexibility prediction method showed the region of 178-184 and 216-222 amino acid residues as a most flexible region (Figure 5:D). On the contrary, Kolaskar & Tongaonkar antigenicity result confirmed the region from 193-212 & 225-240 as highly antigenic (Figure 5:E) while Parker hydrophilicity prediction indicated 217-223 amino acid residues to be more potent (Figure 5:F). Allergenicity pattern of the predicted B-cell epitopes is shown in table 6.

**Table 6:**
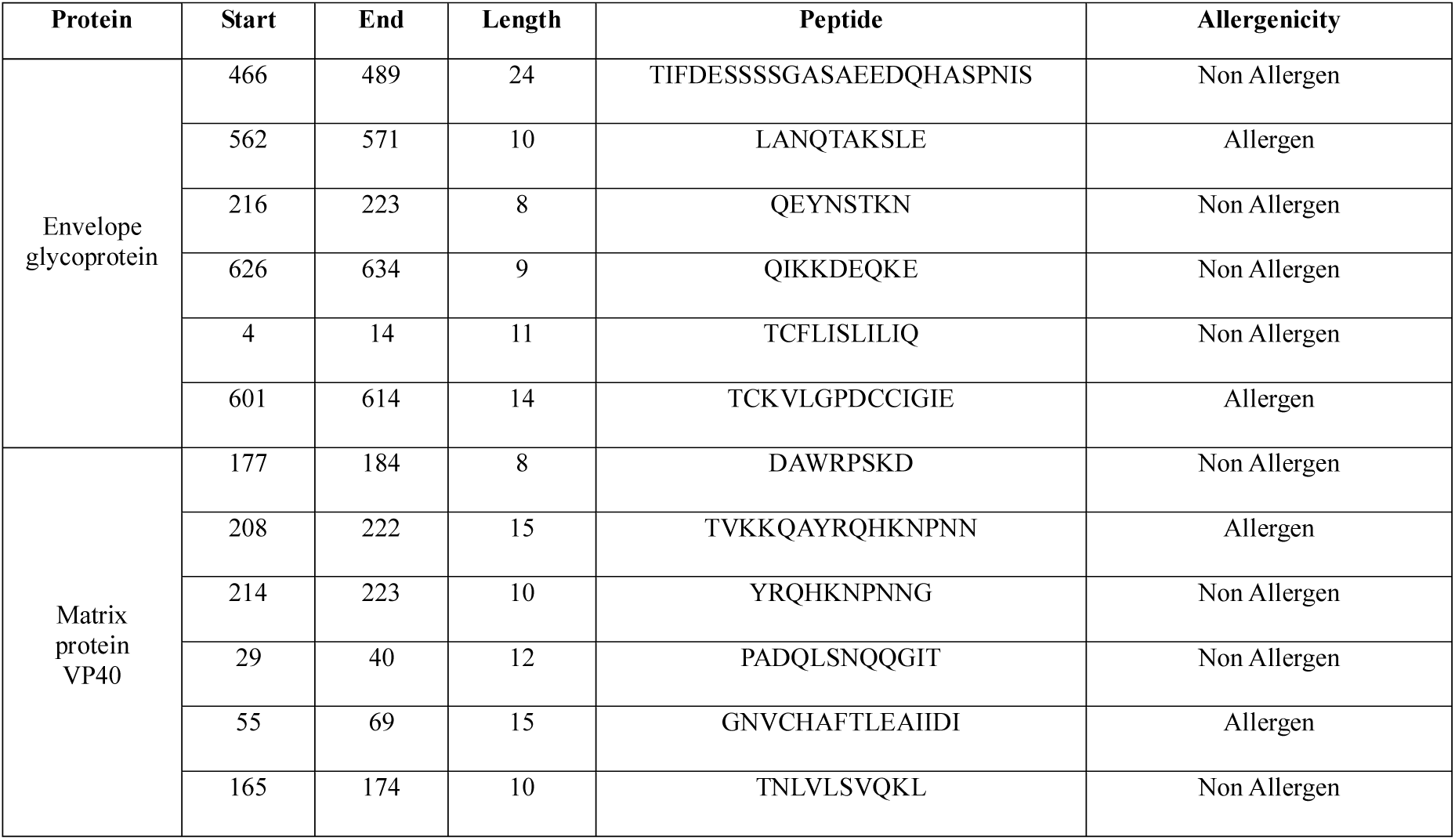
Allergenicity pattern of the predicted B-cell epitopes generated from envelope glycoprotein and matrix protein VP40.

**Fig. 5.**
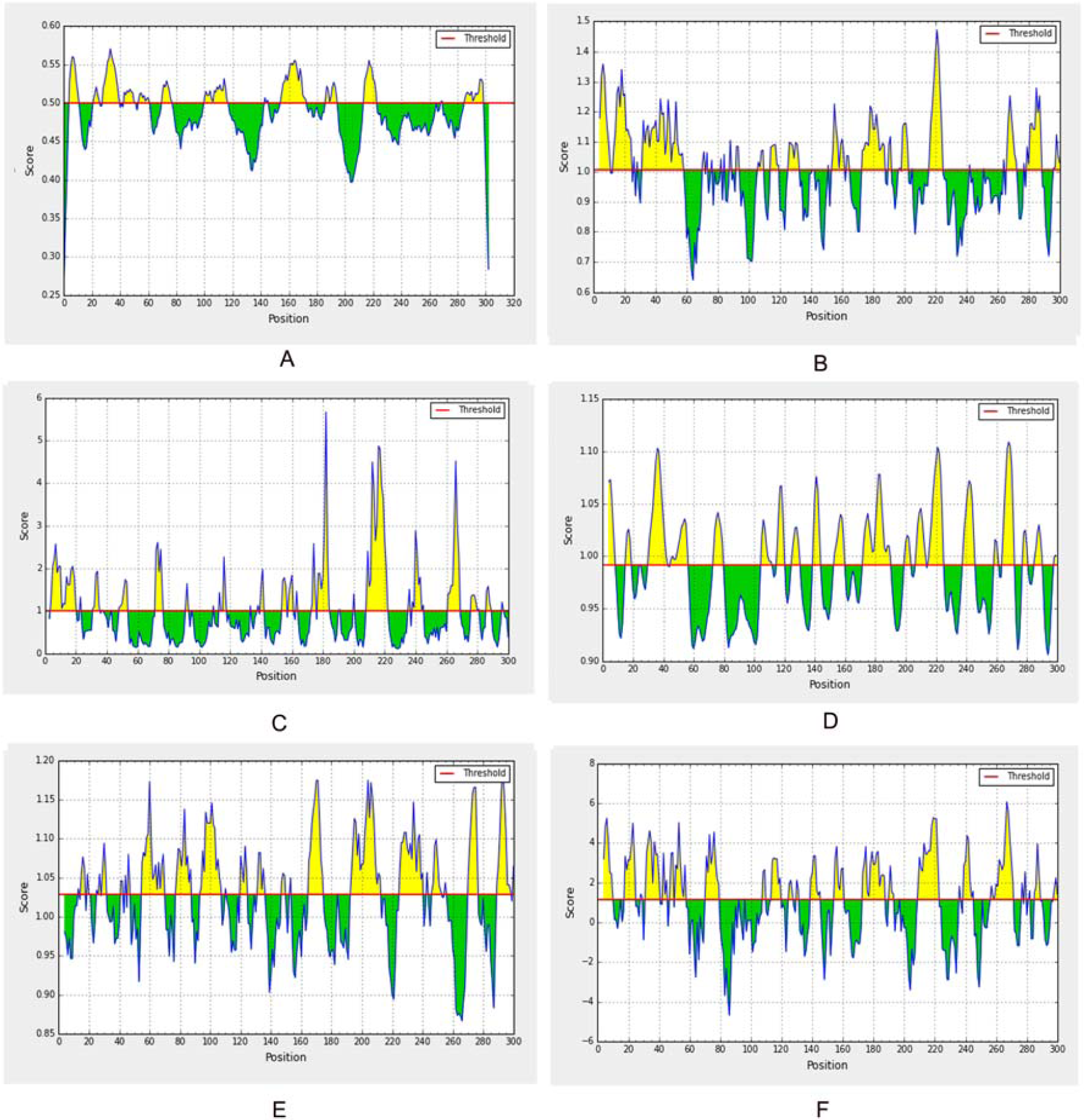
Prediction of B cell linear epitope and intrinsic properties for matrix protein VP40 using different scales (**A:** Bepipred, **B:** Surface accessibility, **C:** Emini surface, **D:** Flexibility, **E:** Antigenicity, **F:** Hydrophilicity). **Notes:** For each graph: x-axis and y-axis represent the position and score; residues that fall above the threshold value are shown in yellow color; the highest peak in yellow color identifies most favored position).

### 3.11. Vaccine construction

Each of the constructs consisted of a protein adjuvant followed by PADRE peptide sequence, while the rest was occupied by the T-cell and B-cell epitopes and their respective linkers. PADRE sequence was incorporated to maximize the efficacy and potency of the peptide vaccine. All three designed vaccines comprised 6 CTL epitopes, 6 HTL epitopes and 8 BCL epitopes. CTL, HTL and BCL epitopes were combined together in a sequential manner using GGGS, GPGPG and KK linkers respectively. The predicted epitopes were separated by suitable linkers to ensure maximal immunity in the body. A total 3 vaccines of 403 (V1), 488 (V2) and 517 (V3) amino acid long were constructed (Table 7) and further analyzed to investigate their immunogenic potential.

**Table 7:**
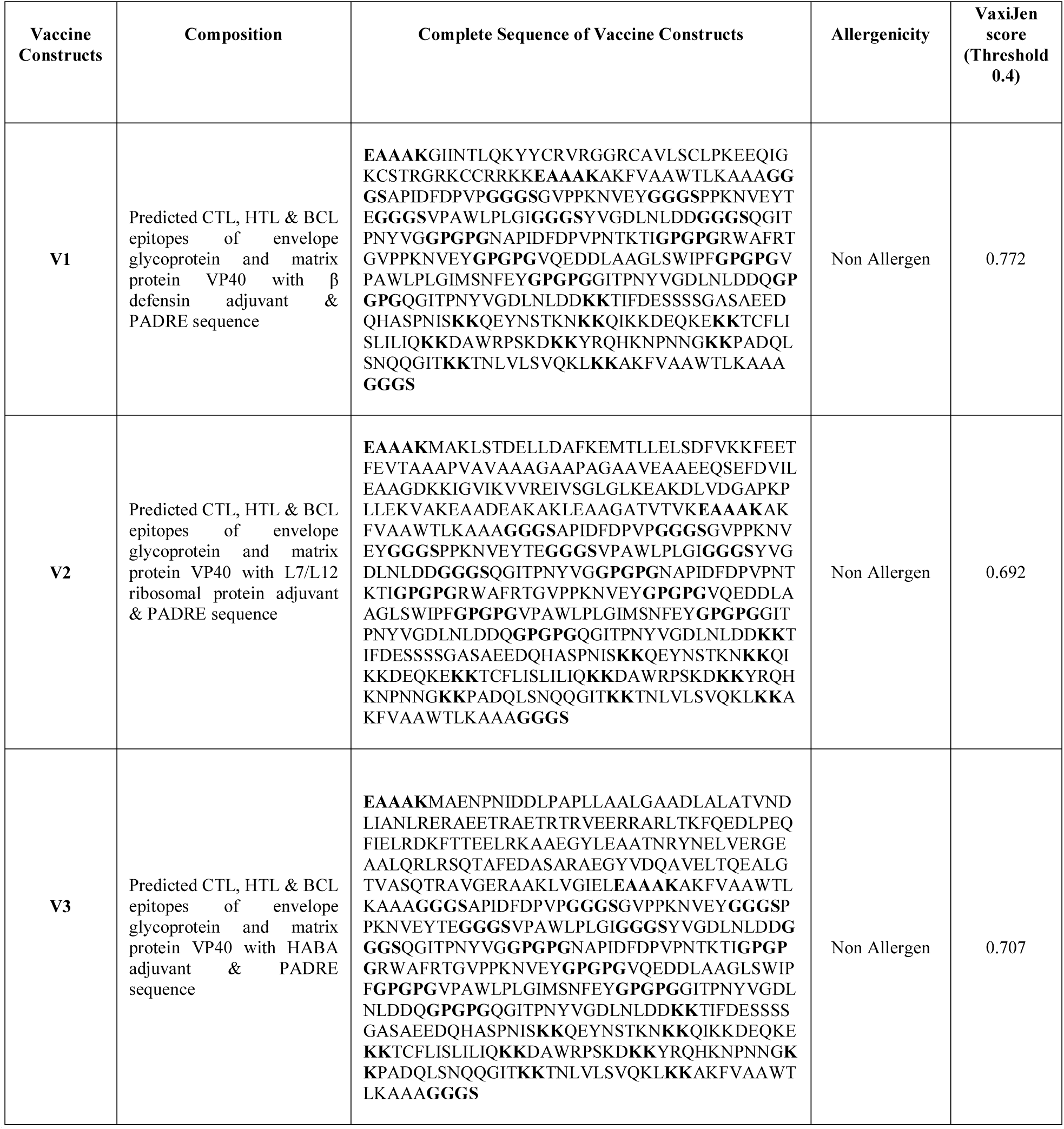
Allergenicity and antigenicity analysis of the constructed vaccines

### 3.12. Allergenicity and antigenicity prediction of different vaccine constructs

AlgPred server was utilized to determine the non-allergic nature of the final vaccine proteins. According to the results, all three constructs (V1, V2 and V3) were non-allergic in behavior. Antigenicity of the vaccine proteins were predicted using VaxiJen 2.0 server. Construct V1 was found superior as potential vaccine candidate with better antigenicity (0.772) and ability to stimulate an efficient immune response (Table 7).

### 3.13. Physicochemical characterization of vaccine protein

The final vaccine construct was characterized on the basis of physical and chemical properties using ProtParam tool. The molecular weight of the vaccine construct was 41.27 kDa which ensures its good antigenic potential. The theoretical pI 9.33 indicating that the protein will have net negative charge above the pI and vice versa. The extinction coefficient was 56380, assuming all cysteine residues are reduced at 0.1% absorption. The estimated half-life of the constructed vaccine was expected to be 1h in mammalian reticulocytes in vitro while more than 10 h in *E. coli* in vivo. Thermostability and hydrophilic nature of the vaccine protein was represented by aliphatic index and GRAVY value which were 65.32 and -0.631 respectively. The computed instability index of the protein was 38.317 which classified it as a stable one.

### 3.14. Secondary and tertiary structure prediction

PSIPRED and NetTurnP 1.0 server were used to predict the secondary structure of the construct V1. The predicted structure of the protein confirmed to have of 21.11% alpha helix, 5.08% sheet and 73.81% coil structure (Figure 06). RaptorX generated the tertiary structure of the designed construct V1 consisting 1 domain only (Figure 7: A and B). The server performed homology modeling by detecting and using 1kj6A from protein data bank (PDB) as best suited template for Vaccine V1. The quality of the 3D model was defined by P value which was 2.63e^-06^ for the predicted vaccine proteint. A low P value ensured better model quality of the predicted vaccine. The Ramachandran plot analysis was also performed to validate the model.

**Fig. 6.**
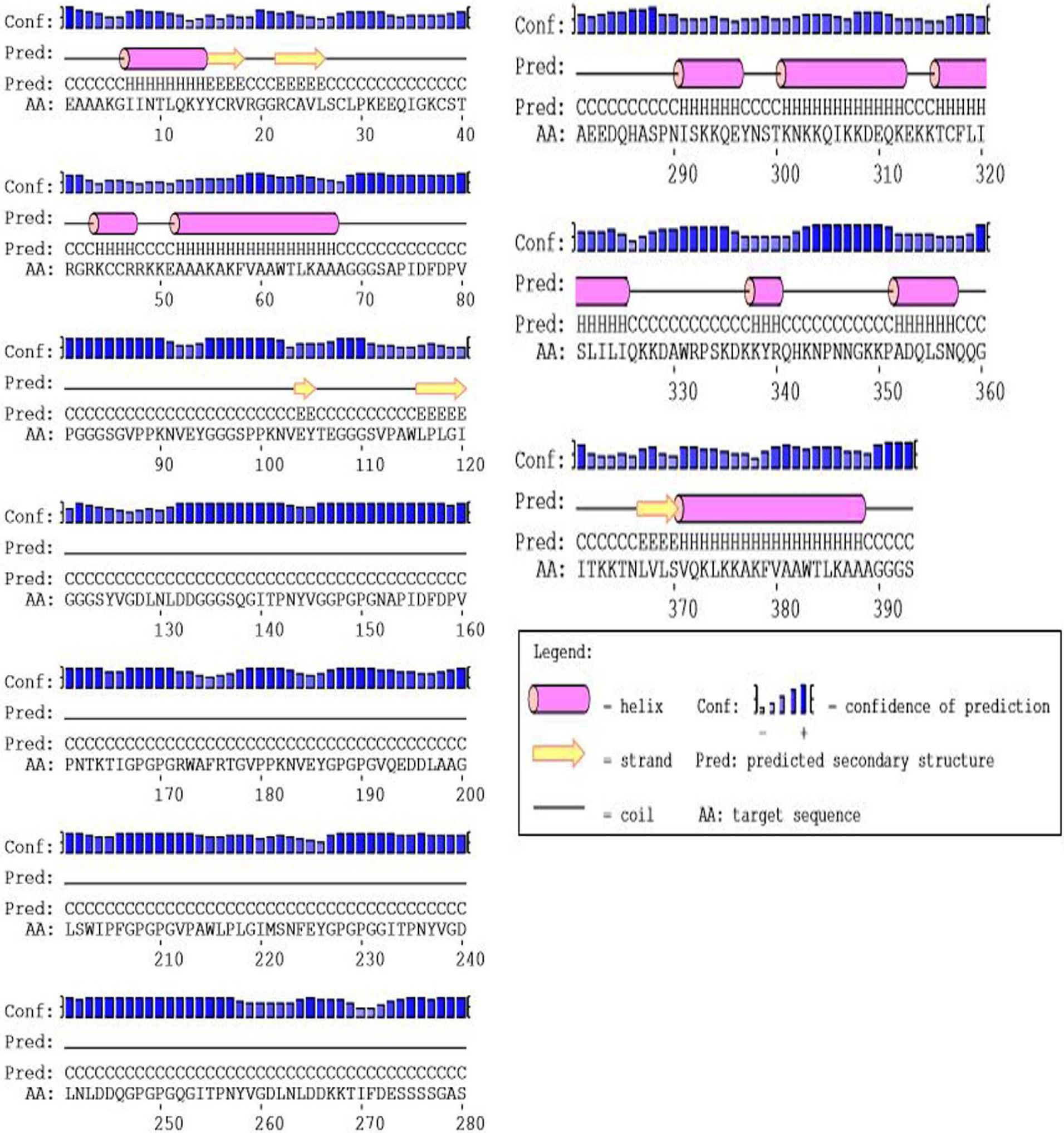
Secondary structure prediction of designed vaccine V1 using PESIPRED server

**Fig. 7.**
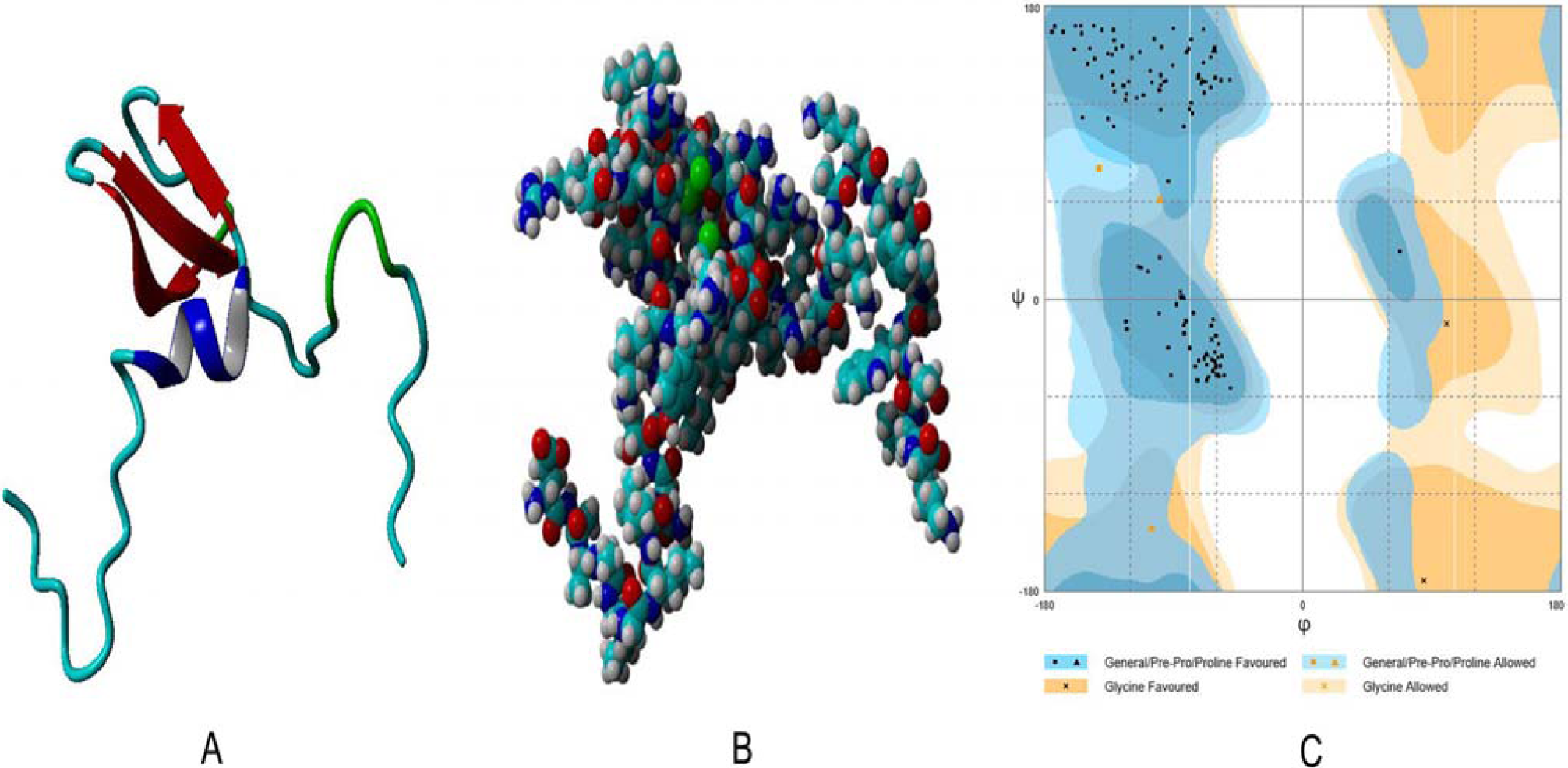
Tertiary structure prediction and validation of vaccine protein V1, **A:** Cartoon format, **B:** Ball structure, **C:** Validation of the 3D structure of vaccine protein V1 by Ramachandran plot analysis.

### 3.15. Tertiary structure refinement and validation

To improve the quality of predicted 3D modeled structure beyond the accuracy, refinement was performed using ModRefiner followed by FG-MD refinement server. During Ramachandran plot analysis 88.9% residues were found in the favored, 9.5% residues in the allowed and 1.6% residues in the outlier region before refinement. However, after refinement 97.5% and 2.5% residues were in the favored and allowed region respectively. No residues existed in the outlier region. (Figure 7:C). Modeled tertiary structure of vaccine construct V2 and V3 have been shown in figure 8.

**Fig. 8.**
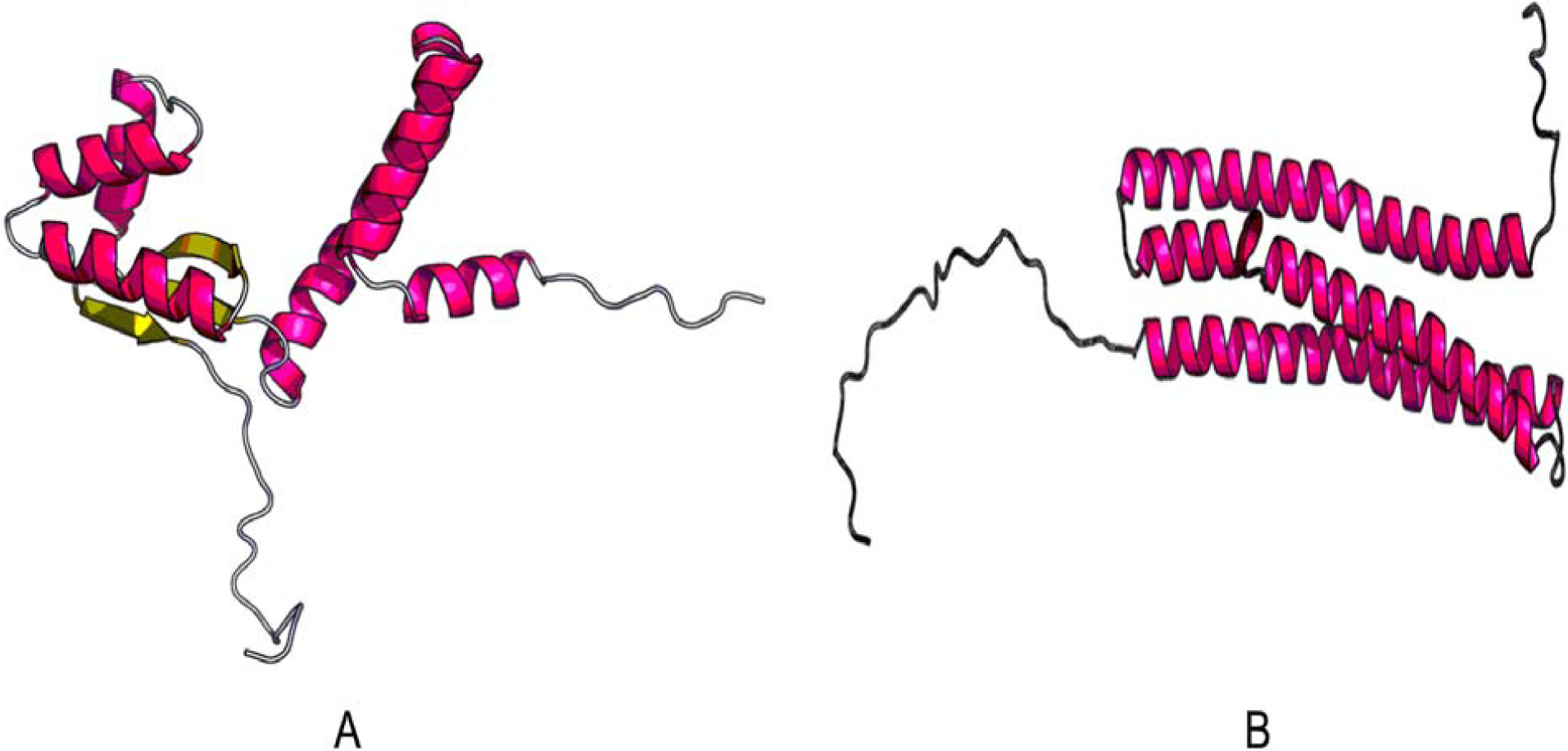
3D modelled structure of vaccine protein V2 **(A)** and V3 **(B)** generated via RaptorX server.

### 3.16. Vaccine protein disulfide engineering

Disulfide engineering was performed by mutating the residues in the highly mobile region of the protein sequence with cystein (Figure 09). A total 17 pairs of amino acid residues were identified with the capacity to form disulfide bond by DbD2 server. After evaluating the residue pairs depending on energy, chi3 and B-factor parameters, only 2 pair satisfied the disulfide bond formation. All 4 residues were replaced with c ALA 4 -VAL 118 and LYS 104 - SER 123 cysteine residue. The value of chi3 considered for the residue screening lay between -87 to +97 while the energy value was less than 2.5.

### 3.17. Protein-protein docking

Again,docking analysis was emloyed between the vaccine constructs and different HLA alleles (Table 8). Construct V1 exhibited biologically significant results and found to be superior in terms of free binding energy. Besides, docking was also undertaken to determine the binding affinity of the designed vaccine construct with human immune TLR8 receptor using ClusPro, HDOC and PatchDock servers (Figure 10) respectively. The ClusPro server generated 30 protein-ligand complexes as output along with respective free binding energy. The lowest energy of -1249.8 was achieved for the complex 9. The hdoc server hypothesized the binding energy for the protein-protein complex was -278.32, while FireDock output refinement of PatchDock server showed the lowest global energy of -29.00 for solution 9.

**Table 8:**
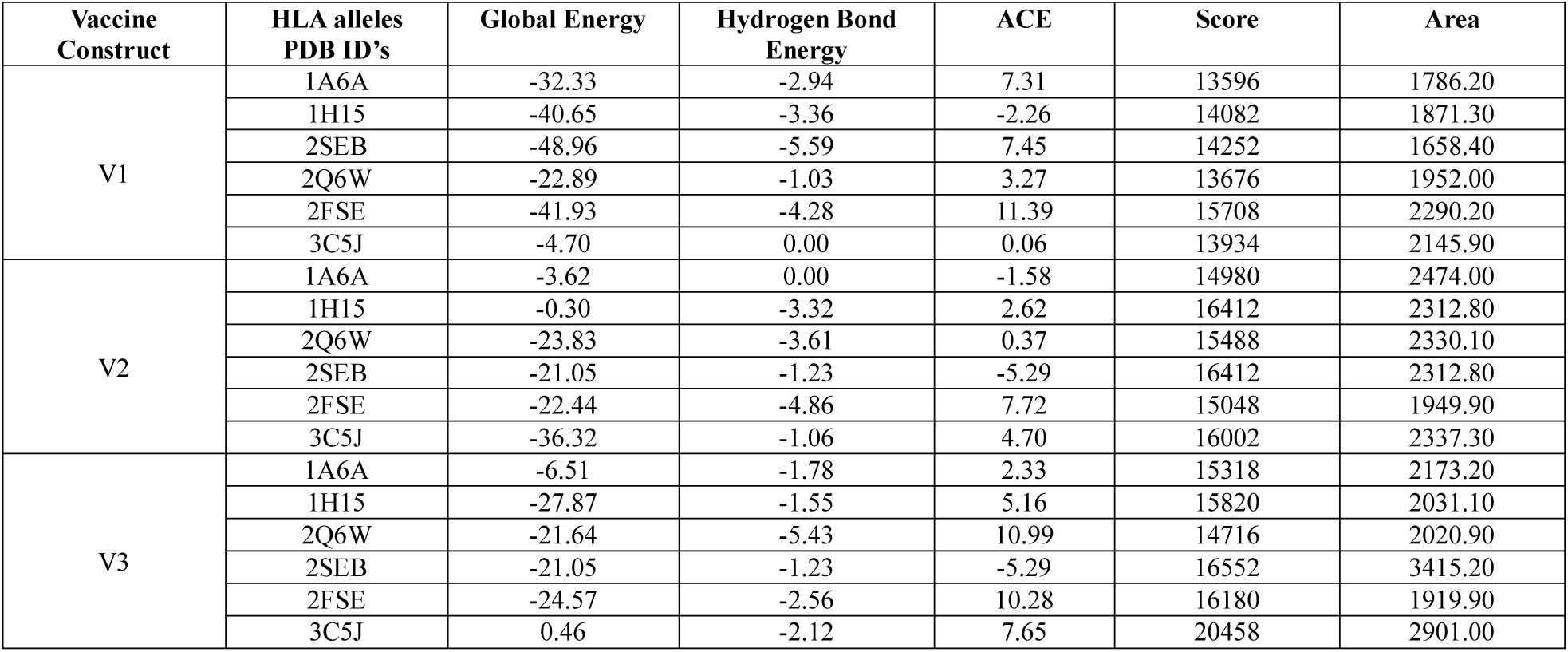
Binding energy of predicted epitopes with selected MHC class I and MHC class II molecules generated from molecular docking by AutoDock

**Fig. 9.**
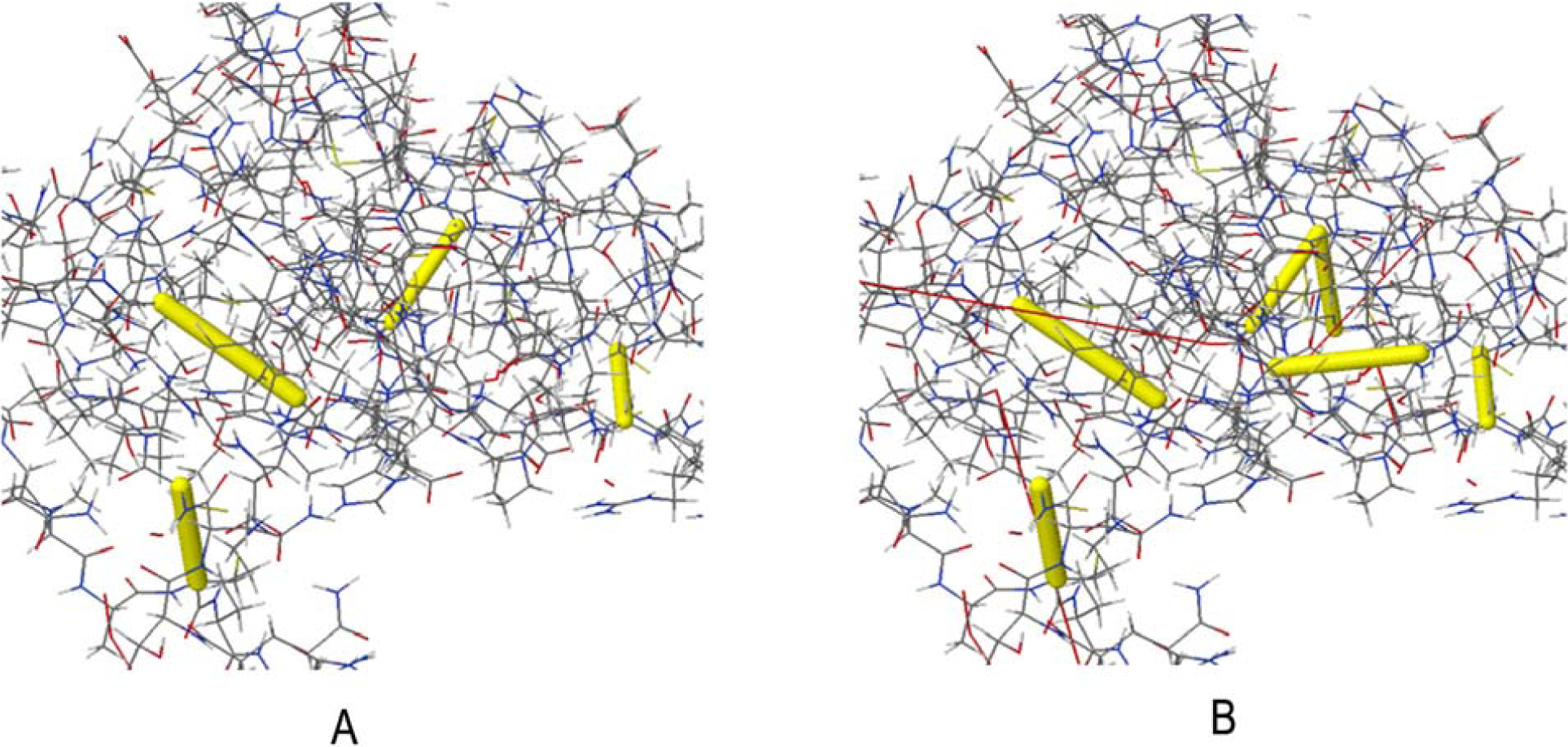
Disulfide engineering of vaccine protein V1; **A:** Initial form, **B:** Mutant form.

**Fig. 10.**
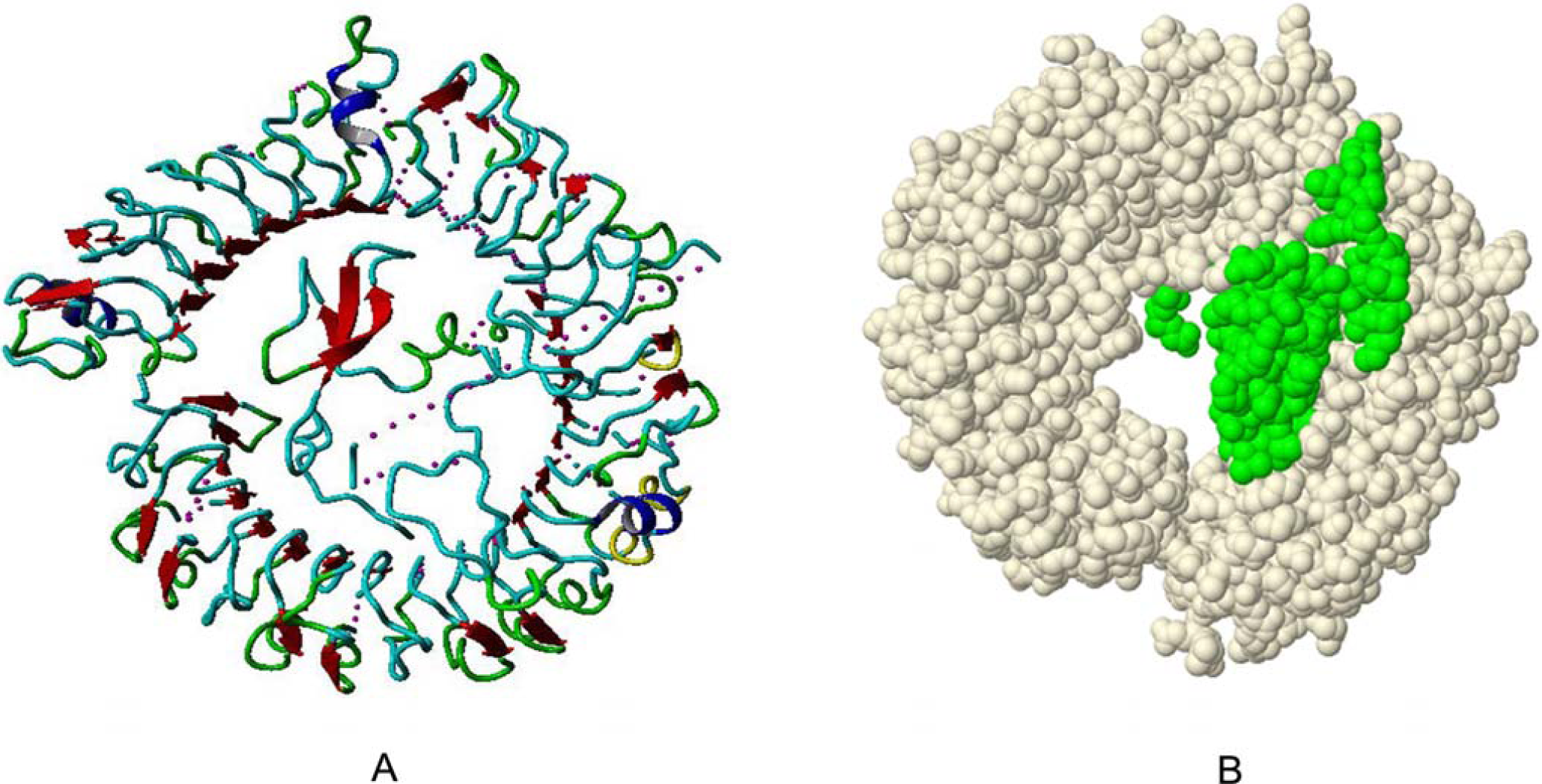
Docked complex of vaccine construct V1 with human TLR8; **A:** Cartoon format and **B:** Ball structure.

**Fig. 11.**
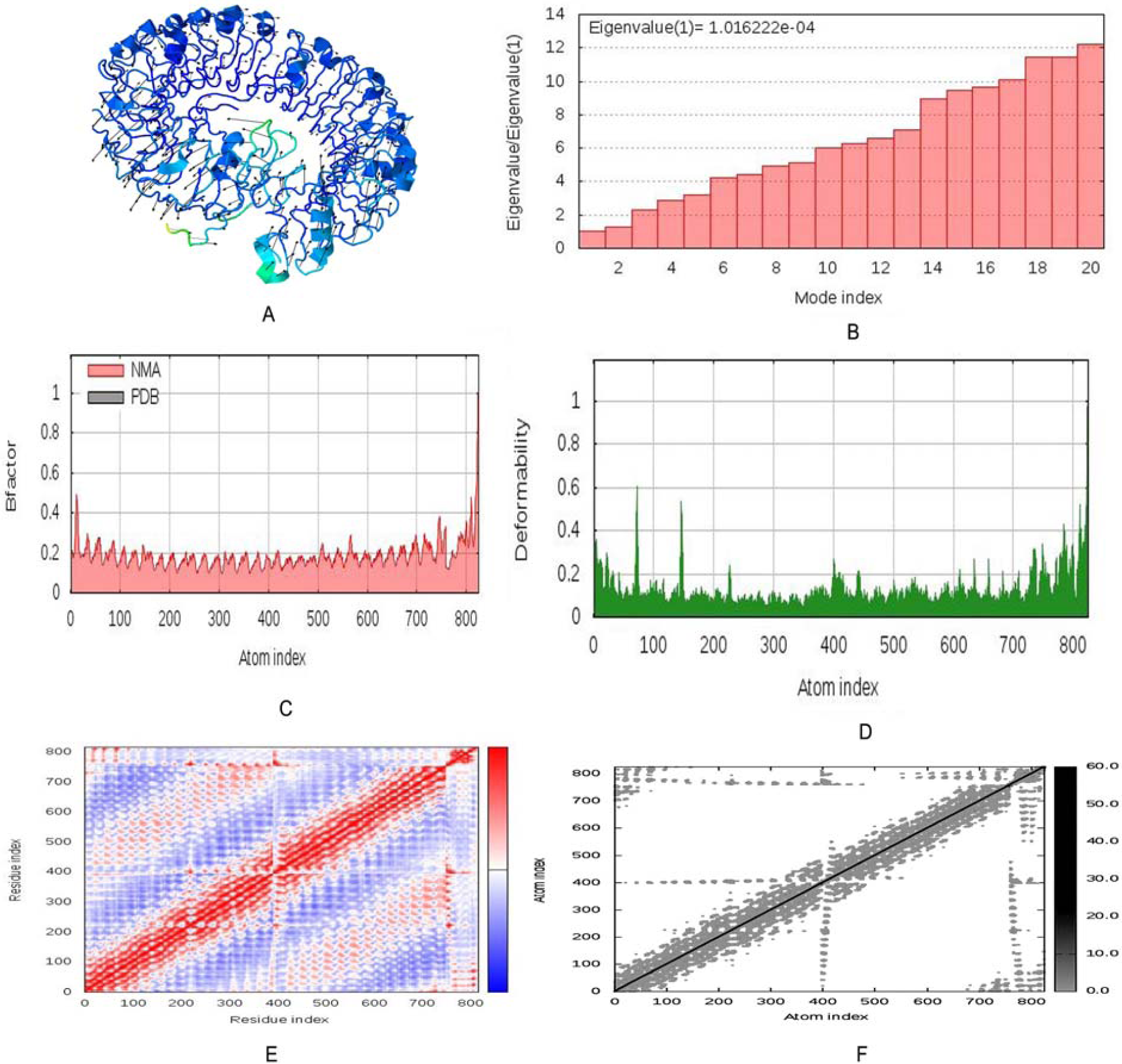
Molecular dynamics simulation of vaccine protein V1-TLR8 complex. Stability of the protein-protein complex was investigated through mobility **(A)**, eigenvalue **(B)**, B-factor **(C)**, deformability **(D)**, covariance **(E)** and elastic network **(F)** analysis.

**Fig. 12.**
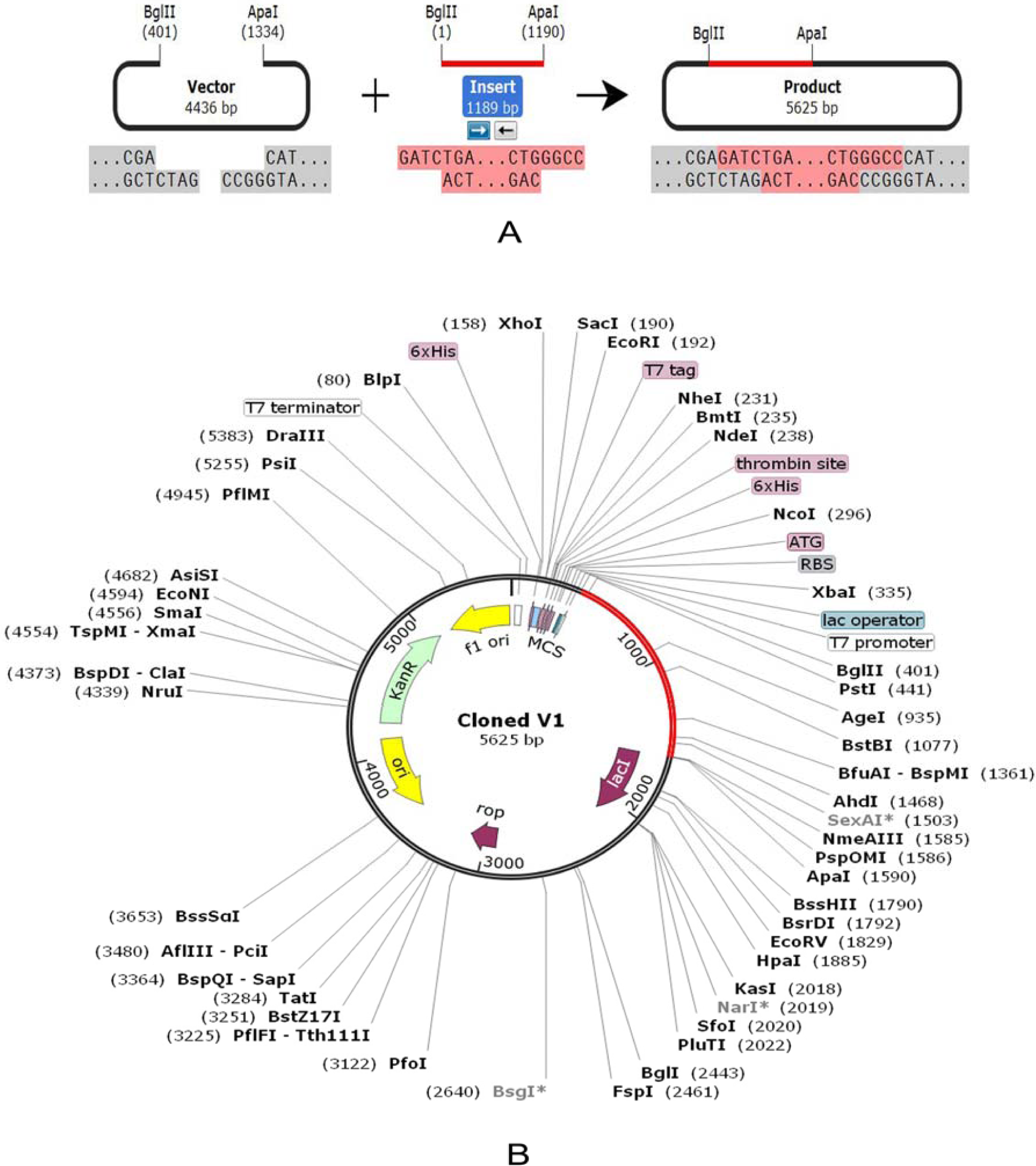
Restriction digestion **(A)** and *in silico* cloning **(B)** of the gene sequence of final vaccine construct V1 into pET28a(+) expression vector. Target sequence was inserted between BglII (401) and ApaI (1334).

### 3.18. Molecular dynamics simulation

Normal mode analysis (NMA) was performed to investigate the stability of proteins and their mobility at large scale. This was rendered by iMODS server by considering the internal coordinates of the docked complex (Fig 14:A). Vaccine protein V1 and TLR were directed towards each other and the direction of each residue in the 3D model was given by arrows. The degree of mobility was indicated by the length of the line. The deformabilty of the complex depends on the individual distortion of each residues, represented by hinges in the chain (Fig 14:D). The eigenvalue found for the complex was 1.0162e^-04^ (Fig 14:B). There was an inverse relationship between eigenvalue and the variance associated to each normal mode.^69^ The B-factor values inferred via NMA, was equivalent to RMS (Fig 14:C). Coupling between pairs of residues was indicated by the covariance matrix where different pairs showed correlated, anti-correlated or uncorrelated motions, represented by red, blue and white colors respectively (Fig 14:E). An elastic network model was also generated which detected the pairs of atoms connected via springs (Fig 14:F). In the diagram each dot was equivalent to one spring between the corresponding pair of atoms and colored according to the level of stiffness. The darker the grays, the stiffer the springs was.

### 3.19. Codon adaptation and *in silico* cloning

Due to dissimilarity in the regulatory systems of human and *E. coli*, codon adaptation was performed considering the expression system of the host. Construct V1 was reverse-transcribed where in the adapted codons, codon adaptation index (CAI) was 0.980 ensuring the higher proportion of most abundant codons. The GC content of the optimized codons (52.671%) was significant as well. The construct did not contain restriction sites for BglII and ApaI and thus indicating its safety for cloning purpose. Finally, the optimized codons were inserted into pET28a(+) vector along with BglII and ApaI restriction sites. A clone of 5625 base pair was produced comprising 1189 bp desired sequence (shown in red color in between the sequence of pET28a(+) vector) and the rest belonging to the vector only (Figure 13).

## 4. Discussion

Marburg virus is known to cause severe hemorrhagic fever in both humans and nonhuman primates with high degree of infectivity and lethality.^1^ To date no approved treatment is available for Marburg virus infection.^2,4^ Therefore, it is essential to take preventive measures against it. The genomic-based technologies continue to transform the field of vaccinology through aiding selection of potential vaccine candidates and facilitating the optimization of the chosen immunogens.

The entire viral proteome of *Marburg marburgvirus* Musoke-80 strain was retrieved from UniProtKB and the physiochemical properties of the proteins were analyzed using ProtParam server. VaxiJen server was used to assess the antigenicity of all the retrieved protein sequences in order to find out the most potent antigenic protein. Among the seven viral proteins, envelope glycoprotein and matrix protein VP40 were identified as the best antigenic protein candidates based on their ability to confer immunity and allowed for further analysis. Vaccine induces production of antibodies that are synthesized by B cells and mediates effector functions by binding specifically to a toxin or a pathogen.^70^ Monovalent vaccines are directed against a specific pathogen or organism that have the ability to trigger both B cell and T cell response.^70^ Cytotoxic CD8+T lymphocytes (CTL) play a vital role by recognizing and killing infected cells or secreting specific antiviral cytokines, thus restricting the spread of infectious agents in the body.^71,72^ Thus, T cell epitope-based vaccination is a unique process of eliciting strong immune response against infectious agents such as viruses.^73^

Approximately 18117 immunogenic epitopes of envelope glycoprotein and 9342 immunogenic epitopes of matrix protein VP40 were generated to be T cell epitopes that can bind a large number of HLA-A and HLA-B alleles with a very high binding affinity using the MHC-I binding predictions of the IEDB with recommended methods. 10800 and 7722 immunogenic epitopes were also generated using MHC-II binding prediction tool of IEDB for envelope glycoprotein and matrix protein VP40 respectively. Top epitopes, bound with the highest number of HLA alleles were selected as putative T cell epitope candidates based on their protein transmembrane topology screening and VaxiJen score.^22^ Today, most vaccines stimulate the immune system into an allergic reaction.^74^ According to the WHO/FAO, if a sequence has an identity of at least six contiguous amino acids over a window of 80 amino acids (0.35% sequence identity) to a known allergens), it is considered to be potentially allergenic. In this study, only the epitopes with non-allergic behavior were allowed for further analysis from both proteins. The result showed that more than 90% population of the world can be covered by the predicted T-cell epitopes. As the MHC super-families play a vital role in vaccine design and drug development, MHC cluster analysis was also performed to determine the functional relationship between MHC variants.

To ensure effective binding between HLA molecules and predicted epitopes, a docking study was performed. The suggested 12 T-cell epitopes from both proteins were subjected to PEP-FOLD3 web-based server for 3D structure conversion. HLA-A*11:01 and HLA-DRB1*04:01 was selected for docking analysis with MHC class I and class II binding epitopes respectively. Epitope ‘VQEDDLAAGLSWIPF’ from envelope glycoprotein was found to be best as it bound in the groove of the HLA-DRB1*04:01 with an energy of - 7.8 kcal/mol with significant numbers of hydrogen bonding. Again, VP40-epitope ‘VPAWLPLGIMSNFEY’ containing the 9-mer core ‘VPAWLPLGI’ was found best in terms of binding energy (-7.0 Kcal/mol). However, in this study we developed a multi-epitope subunit vaccine to ensure better immune protection. All the finalized epitopes showed a lower binding energy which was biologically significant.

For B-cell epitope prediction, we predicted amino acid scale-based methods for the identification of potential B-cell epitopes using four algorithms from IEDB server. The most potent B cell epitopes for envelope glycoprotein and matrix protein VP40 were identified as vaccine candidates against Marburgvirus. The final vaccine proteins were constructed using the promiscuous epitopes and protein adjuvants along with PADRE sequence. Individual epitopes were linked together via suitable linker to ensure effective immune response. Literature studies revealed that PADRE containing vaccine construct showed better CTL responses than the vaccines lacked it.^75^

The constructed vaccines were further checked for their non-allergic behavior and immunogenic potential. Construct V1 was superior in terms of antigenicity and Vaxigen score. The physicochemical properties and secondary structure of V1 was also analyzed before tertiary structure prediction and refinement of 3D model. To strengthen our prediction, we checked the interaction between our vaccine construct with different HLA molecules (DRB1*0101, DRB3*0202, DRB5*0101, DRB3*0101, DRB1*0401, and DRB1*0301). Again construct V1 was found to be best considering the free binding energy. Moreover, docking analysis was also performed to explore the binding affinity of vaccine protein V1 and human TLR8 receptor to evaluate the efficacy of used adjuvant. Molecular dynamics study was conducted to determine the complex stability as well. Structural dynamics had been investigated previously using subsets of atoms and covariance analysis^60^ Literature studies linked the stability of macromolecules with correlated fluctuations of atoms.^76,77^ Essential dynamics was compared to the normal modes of proteins to determine its stability through iMODS server. The analysis revealed negligible chance of deformability for each individual residues, as location of hinges in the chain was not significant and thereby strengthening our prediction. Finally, the designed vaccine construct V1 was reverse transcribed and adapted for *E. coli* strain K12 prior to insertion within pET28a(+) vector for its heterologous cloning and expression. Our predicted *in silico* results were based on different analysis of sequence and various immune databases. We suggest further wet lab based analysis using model animals for experimental validation of the predicted vaccine candidates.

## 5. Conclusion

Prevention of newly emerging Marburg virus infections and control during outbreaks is both mandatory and challenging. *In silico* studies can guide the experimental work for finding the desired solutions with fewer trials and error repeats and thus saving both time and costs for the researchers. A novel chimeric subunit vaccine (V1) against Marburgvirus was developed using the most epitopes antigenic via various bioinformatics tools. Thus our study provides new and valuable epitope candidates and prompts the future vaccine development against Marburgvirus as well as other infectious diseases. However, we suggest for further wet lab based experiments for the acceptance and validation of the modelled vaccine.

## Abbreviations

MHF: Marburg hemorrhagic fever
MARV: Marburg virus disease
NCBI: National center for Biotechnology information
IEDB: Immune epitope database and analysis resource
CTLs: Cytotoxic T lymphocytes
MGL: Molecular graphics laboratory
MHC: Major histocompatibility complex

### Acknowledgments

Authors like to acknowledge the authority of the Bioinformatics Laboratory of Sylhet Agricultural University for the technical support of the project.

## Author contributions

Mahmudul Hasan: Conceptualization, Supervision, Project administration and reviewing. Kazi Faizul Azim: Experiment Design, Data Handling, Data Analysis, Manuscript writing and draft Preparation

Aklima Begum, Noushin Anika Khan, Tasfia Saiyara Shammi: Data Handling and Data analysis

Md. Abdus Shukur Imran, Ishtiak Malique Chowdhury: Data Handling

Shah Rucksana Akhter Urme: Data Analysis

